# Mature interneuron subtypes arise from distinct spatial and temporal subdomains within the caudal ganglionic eminence

**DOI:** 10.1101/2025.07.28.667082

**Authors:** Vedant Garg, Samra Beyene, Anthony Tanzillo, Allison Tucker, Aidan Johantges, Daniel Abebe, Yajun Zhang, Timothy J. Petros

## Abstract

Forebrain inhibitory interneurons are born from transient structures during embryogenesis known as the medial and caudal ganglionic eminences (MGE and CGE, respectively). The MGE and CGE generate distinct, non-overlapping cohorts of interneurons that can be defined by their transcriptomic, morphological, and electrophysiological characteristics. In the MGE, somatostatin-expressing (SST+) cells arise predominantly from the dorsal-posterior MGE from E12-E16 whereas parvalbumin-expressing (PV+) cells are born in the ventral-anterior MGE throughout embryogenesis. This relationship between spatiotemporal origin and mature interneuron subtypes has led to genetic insights regarding fate and maturation of these MGE-derived cells. A similar organization has never been explored in the CGE, despite the significant increase in CGE-derived interneurons in primates compared to rodents. Here we harvested fluorescent cells from distinct CGE subdomains at E13.5 and E15.5 and grafted them into WT neonatal mice cortices. One month post-transplantation, brains were immunostained for interneuron markers to relate mature CGE-derived interneurons with spatiotemporal origins within the CGE. Our results indicate that there are significant spatial biases in the CGE, with specific interneuron subtypes preferentially arising from distinct CGE subdomains. These biases are relatively stable over time, implying a minimal relationship between temporal birthdate and interneuron subtype. In the future, combining these insights with spatial transcriptome profiles will generate critical insights into gene regulation of CGE-derived interneurons.

## INTRODUCTION

The mammalian cortex is composed of excitatory glutamatergic projection neurons and inhibitory GABAergic interneurons that generate the intricate excitatory-inhibitory balance required for normal brain function. Interneurons are an incredibly heterogeneous cell population, and perturbation of their maturation and function is associated with diseases such as epilepsy, autism spectrum disorders, schizophrenia, and numerous other neurodevelopmental and psychiatric disorders (1–4). Notably, many genes implicated in these brain disorders are strongly enriched in neural progenitors, and in some instances specifically in developing interneurons (5–9), which emphasizes the need to understand mechanisms regulating interneuron fate and maturation.

Forebrain interneurons are generated in transient structures in the ventral telencephalon known as the ganglionic eminences (GE), specifically the medial, caudal, and lateral ganglionic eminences (MGE, CGE, and LGE, respectively). Transplantation studies provided the first critical evidence that cells originating from the MGE, CGE and LGE have different migratory behaviors, cortical layering properties, and cell fates (10–15). These studies revealed that grafted interneuron progenitors can migrate extensively throughout the cortex, differentiate into interneuron subtypes with appropriate laminar organization, integrate into local circuitry, and exhibit similar electrophysiological properties to those of their endogenous counterparts. More refined transplantation studies focused specifically on the MGE revealed both a spatial and temporal logic relating MGE subdomains with mature interneuron fate: SST+ interneurons arise primarily from the dorsal-caudal MGE at earlier embryonic ages (E12.5-E15.5) whereas PV+ interneurons have a complimentary bias from the ventral-rostral MGE and are born throughout neurogenesis (E12.5-E18.5) (16–20). Combining this spatial origin-interneuron subtype relationship with transcriptional profiles permitted greater in-depth analysis of genetic regulation of cell fate, such as the dorsal MGE enriched gene *Nkx6.2* driving formation of SST+/CR+ interneurons (17, 21, 22). Additionally, work over the last several decades has uncovered factors that regulate initial fate decisions within the MGE, including spatial gradients of morphogens (specifically Shh and Wnts), temporal cell birthdates, and the mode of neurogenesis (23–25).

However, our understanding of mechanisms regulating specification of CGE-derived interneuron subtypes is largely unknown. While the transcription factor *Nkx2.1* acts as a ‘master regulator’ for MGE development (26), a similar genetic driver of CGE fate has not been identified. Morphogen gradients that pattern the CGE are poorly understood, and a relationship between spatiotemporal origins and mature CGE-derived interneuron subtypes has not been explored. While CGE-derived interneurons make up ∼30% of cortical interneurons in the mouse (27), this population is significantly increased in humans and non-human primates (28–33), with the largest increase in CGE-derived VIP+ disinhibitory bipolar interneurons (34, 35). There is evidence that CGE-derived interneurons may be a critical nexus for multiple neuropsychiatric disorders (36–38), and the CGE is the primary source of tumors and lesions in tuberous sclerosis complex (TSC) (39). Thus, a better understanding of mechanisms regulating fate and maturation of CGE-derived interneurons is needed to understand both normal brain development and potential disease etiologies.

Here we transplanted fluorescent CGE cells into the cortices of WT pups to determine if specific CGE-derived interneuron subtypes preferentially arise from distinct spatiotemporal domains within the CGE. We find that some mature interneuron subtypes showed strong spatial biases, arising primarily from one subdomain within the CGE, while other subtypes were generated relatively homogenously throughout the CGE. These spatial biases were relatively consistent at E13.5 and E15.5, indicating that temporal birthdates of CGE-derived cells are not strongly correlated with specific interneuron subtypes. This defined spatiotemporal logic characterizing the origins of CGE-derived interneurons, in combination with differential gene expression patterns in the CGE (40), paves the way to understand initial fate decisions of this important yet understudied cohort of forebrain interneurons.

## RESULTS

### Tomato+ transplanted CGE cells migrate throughout the cortex and predominantly settle in superficial cortical layers

To identify spatial and temporal origin biases of CGE-derived interneuron subtypes, we dissected the anterior, posterior, ventral and dorsal CGE subdomains (aCGE, pCGE, vCGE and dCGE, respectively) from embryonic mice and transplanted them into WT neonatal pups. As there is not a CGE-specific Cre-driver, we generated *Dlx5/6-Cre;Ai9* mice, which labels all postmitotic MGE, LGE and CGE cells with tdTomato (Tom+). Since the LGE and CGE are anatomically continuous, we defined the posterior 50% of the LGE-CGE structure as the CGE. To harvest aCGE and pCGE, the CGE was divided into thirds along the antero-posterior axis, and the most anterior and posterior thirds were collected. To harvest vCGE and dCGE, the CGE was bisected longitudinally, and the dorsal and ventral halves were collected (Figure 1A).

**Figure 1.**
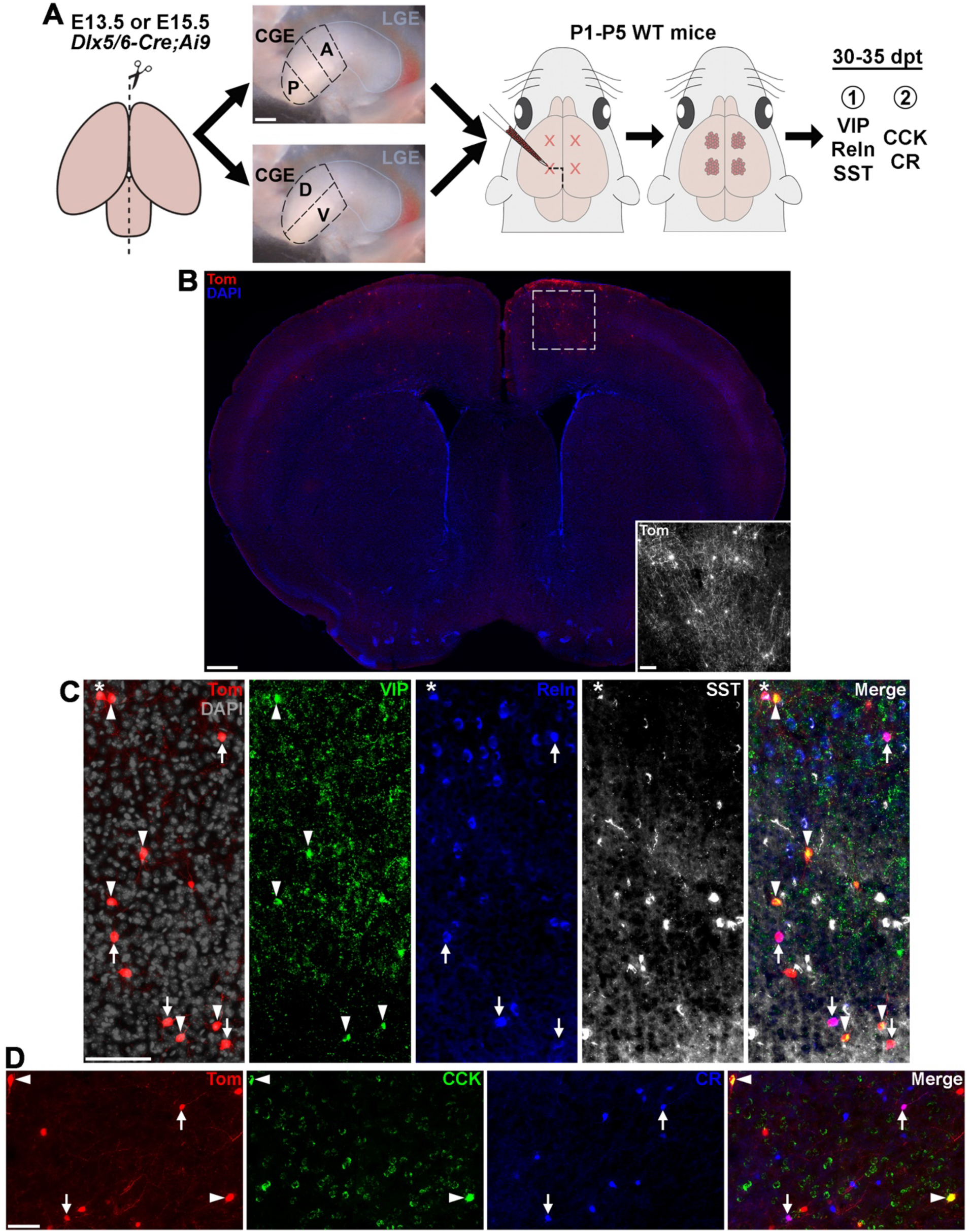
Transplanted CGE cells express markers of mature CGE-derived interneuron subtypes. **A.** Schematic depicting cell transplantation procedure of CGE subdomains from E13.5 and E15.5 *Dlx5/6-Cre;Ai9* mice into cortex of WT neonate pups. Scale bar = 200 μm. **B.** Brain section ∼35 days post transplantation depicting tdTom+ cells dispersed throughout the cortex in both hemispheres. Scale bar = 500 μm; inset = 100 μm. **C.** Example of transplanted CGE-derived Tom+ cells expressing VIP (arrowheads) or Reln (arrows), and MGE-derived Reln+/SST+ (asterisk) cells. Scale bars = 50 μm. **D.** Example of transplanted CGE-derived Tom+ cells expressing CCK (arrowheads) and CR (arrows). Scale bar = 50 μm.

We performed dissections at E13.5 and E15.5 to capture two different timepoints representing early and peak CGE neurogenesis (41). While CGE cells are generated until birth (41), the transition from ganglionic eminences to more mature structures combined with loss of clear GE boundaries presents technical challenges for cleanly dissecting distinct CGE subdomains at later embryonic ages. Single cell dissociations were obtained (42) and transplanted into the cortices of P1-P5 WT pups as previously described (43, 44). Brains were collected 30-35 days post transplantation and stained for markers of mature CGE-derived interneuron subtypes. Tom+ cells were clearly visible in the host brains and migrated large distances throughout the cortex (Figure 1B). While MGE-derived interneurons are located broadly across all cortical layers, CGE-derived interneurons show a strong bias towards superficial cortical layers due (41, 45, 46). Dividing the cortex into 10 equidistant bins reveals that grafted Tom+ CGE-derived cells follow this same pattern (Figure S1A). On average, CGE-derived cells are located 2x more superficial compared to grafted MGE cells, with strongest enrichment in the most superficial bin that primarily represents layer I (∼40% of CGE-derived cells located in bin 1 vs. ∼14% of MGE cells) (Figure S1B-C). Conversely, MGE-derived cells were relatively evenly distributed throughout cortical layers.

### Tomato+ transplanted CGE cells express markers of mature CGE-derived interneuron subtypes

MGE-derived interneurons can be segregated into largely non-overlapping parvalbumin-expressing (PV+) fast-spiking and somatostatin-expressing (SST+) non-fast spiking cardinal classes, which can then be further parsed into additional PV+ and SST+ subtypes (47–49) (Figure S2A-B). The CGE lacks a clean classification scheme because many traditional markers for CGE-derived classes are expressed by other cell types or partially overlap with each other, which is comprehensively reviewed in Machold & Rudy (50). For example, cholecystokinin-expressing (CCK+) basket cells are a well-characterized subset of CGE-derived cells, but CCK mRNA is also expressed by some PV+ MGE-derived interneurons and excitatory cells in the cortex and hippocampus (47, 51). The vasoactive intestinal peptide-expressing (VIP+) class of interneurons are disinhibitory and preferentially synapse on SST+ interneurons, but subsets of VIP+ cells co-express CCK or another interneuron marker calretinin (CR, encoded by *Calb2*) (50). The two best complimentary markers that label nearly all CGE-derived interneurons are VIP and Reelin (Reln), but many MGE-derived SST+ cells also express Reln (Figure S2C). Other markers such as Lamp5, Ndnf and Sncg have more recently been shown to label specific CGE-derived subtypes with varying levels of specificity (50).

We chose 2 staining conditions that form complimentary pairs and cover the breadth of mature CGE-derived subtypes. One cohort of brain sections were stained for VIP, Reln, and SST (Figure 1C). The Tom+/Reln+/SST+ cells are MGE-derived interneurons known to migrate through the CGE which will ‘contaminate’ CGE dissections in these *Dlx5/6-Cre;Ai9* mice (15). By subtracting this population from all Reln+ cells, we can analyze the largely complementary VIP+ and Reln+ CGE-derived interneuron population (Figure S2C). Another cohort of brain sections was stained for CCK and CR (Figure 1D). CR+ CGE cells label a subpopulation of VIP+ cells, as well as a population of Reln+ CGE cells. This CR+ population is largely distinct from the CCK+ basket cells, which also partially overlap with VIP+ and Reln+ cells (Figure S2C). While CCK mRNA is observed in mature PV+ MGE-derived cells (Figure S2C), we do not observe CCK protein expression in PV+ cells in the cortex or hippocampus (Figure S3). Thus, we are confident that the Tom+/CCK+ are CGE-derived putative basket cells and not MGE-derived PV+ basket cells. These CCK+ and CR+ cohort of cells are largely complimentary and cover a large swath of CGE-derived interneurons.

We characterized cell types from 99 mice that received Tom+ cell transplants for the 8 different conditions (4 brain regions from 2 timepoints each), with a range of 9-14 brains per condition and an average of 410 Tom+ cells counted per WT host brain. For all conditions, an average of 64.8% Tom+ cells could be classified into one of the CGE-derived interneuron subtypes (CCK+, CR+, VIP+, Reln+/SST-), with another 10.6% of Tom+ cells representing MGE-derived SST+ cells migrating through the CGE (Figures 2A-B & S4). The remaining ∼25% of Tom+ cells that were unlabeled with interneuron markers is consistent with previous results for transplanted MGE-derived cells (43) and likely represent cells that did not mature properly after transplantation, or possibly are GABAergic projection neurons that originate from the CGE and do not express these interneuron markers. All statistics comparing temporal differences in cell types between E13.5 and E15.5, and spatial differences between distinct CGE subdomains at E13.5 and E15.5, are presented in Supplementary Tables 1-3. As detailed below, we observed significant spatial differences in the CGE, with many cell types showing strong origin biases from distinct CGE subdomains. Conversely, we observed minimal differences in subtype origin over time, with the spatial biases largely consistent between E13.5 and E15.5.

**Figure 2.**
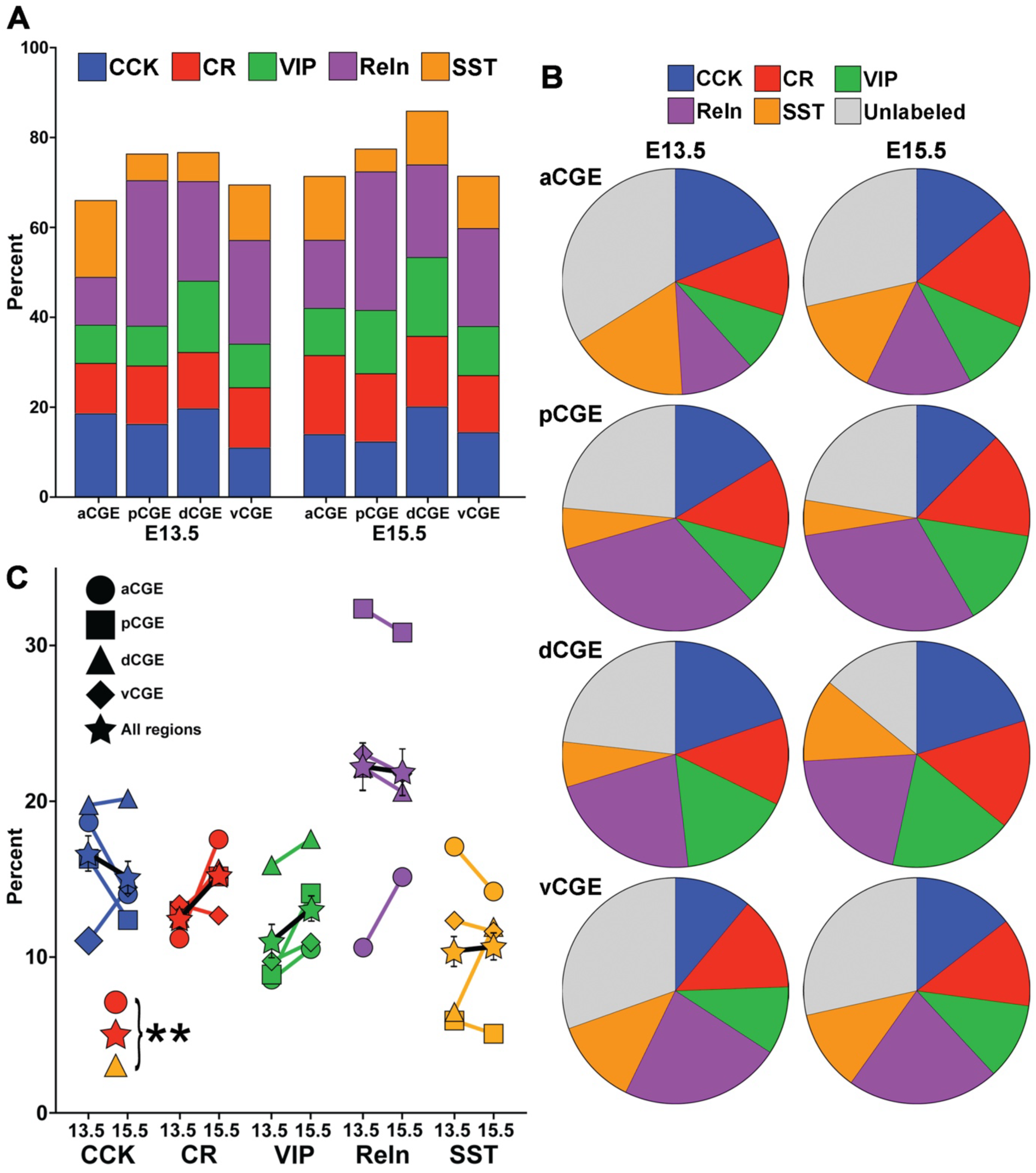
Proportion of interneuron subtypes arising from CGE subdomains is relatively consistent from E13.5 to E15.5. **A.** Bar graph displaying proportion of grafted Tom+ cells that express CCK, CR, VIP, Reln (Reln+/SST-) and SST (Reln+/SST+). **B.** Pie charts depicting proportion of Tom+ cells expressing each marker for all 8 experimental conditions. **C.** Line graphs depicting average percent of Tom+ cells expressing each marker at E13.5 and E15.5 for each CGE subdomain, as well as all subdomains combined (star). Data points not shown, and only s.e.m. bar depicted for combined subdomains for visualization purposes. Three conditions have a significant increase in proportional expression from E13.5 to E15.5: CR aCGE (red circle), CR combined regions, (red star), SST dCGE (yellow triangle). No other conditions reached significance. ** p < 0.01.

### Origin of CGE-derived subtypes is relatively constant from E13.5 to E15.5

Comparing E13.5 vs. E15.5 revealed that origins of CGE-derived interneurons are relatively stable over time. The biggest difference was with CR+ cells, with a significant increase in the overall number of CR+ cells from the E15.5 transplants, and specifically within the aCGE region, compared to E13.5 (Figure 2C & Supplementary Table 1). We also observed a trend for increased CR+ cells from the pCGE and dCGE (Supplementary Table 1) at E15.5, indicating that there is likely increased production of CR+ interneurons within the CGE at E15.5 compared to E13.5. The only other datapoint reaching significance was an increase in SST+ cells in the dCGE at E15.5 compared to E13.5, which likely arises from increased MGE-derived SST+ cells migrating through this region at later embryonic ages. There was a greater percentage of VIP+ cells arising from all CGE regions at E15.5 compared to E13.5, but none of these datapoints reached significance.

### Interneuron subtypes display significant biases for originating from distinct CGE subdomains

We identified strong spatial biases for most CGE-derived interneuron subtypes within the CGE. Analysis of variance (ANOVA) for all 5 cell types at E13.5 and E15.5 (10 conditions) revealed a significant difference of cell type origin within CGE subdomains in 8 of 10 conditions, and a ninth condition (E15.5 CCK) having a P-value = 0.051 (Supplementary Tables 2-3). CR+ and CCK+ cells had more moderate spatial biases compared to other cell types. CR+ cells from E13.5 transplants showed no spatial biases, and there was only a significant difference between aCGE and vCGE for CR+ cells from E15.5 transplants (Figure 3A). Thus, CR+ cells, which showed the greatest temporal bias, display minimal spatial bias within the CGE. CCK+ cells also displayed moderate spatial biases, with dCGE transplants having the highest percentage of CCK+ cells at both ages, reaching significance compared to other brain regions at both E13.5 (vs. vCGE) and E15.5 (vs. pCGE) (Figure 3A).

**Figure 3.**
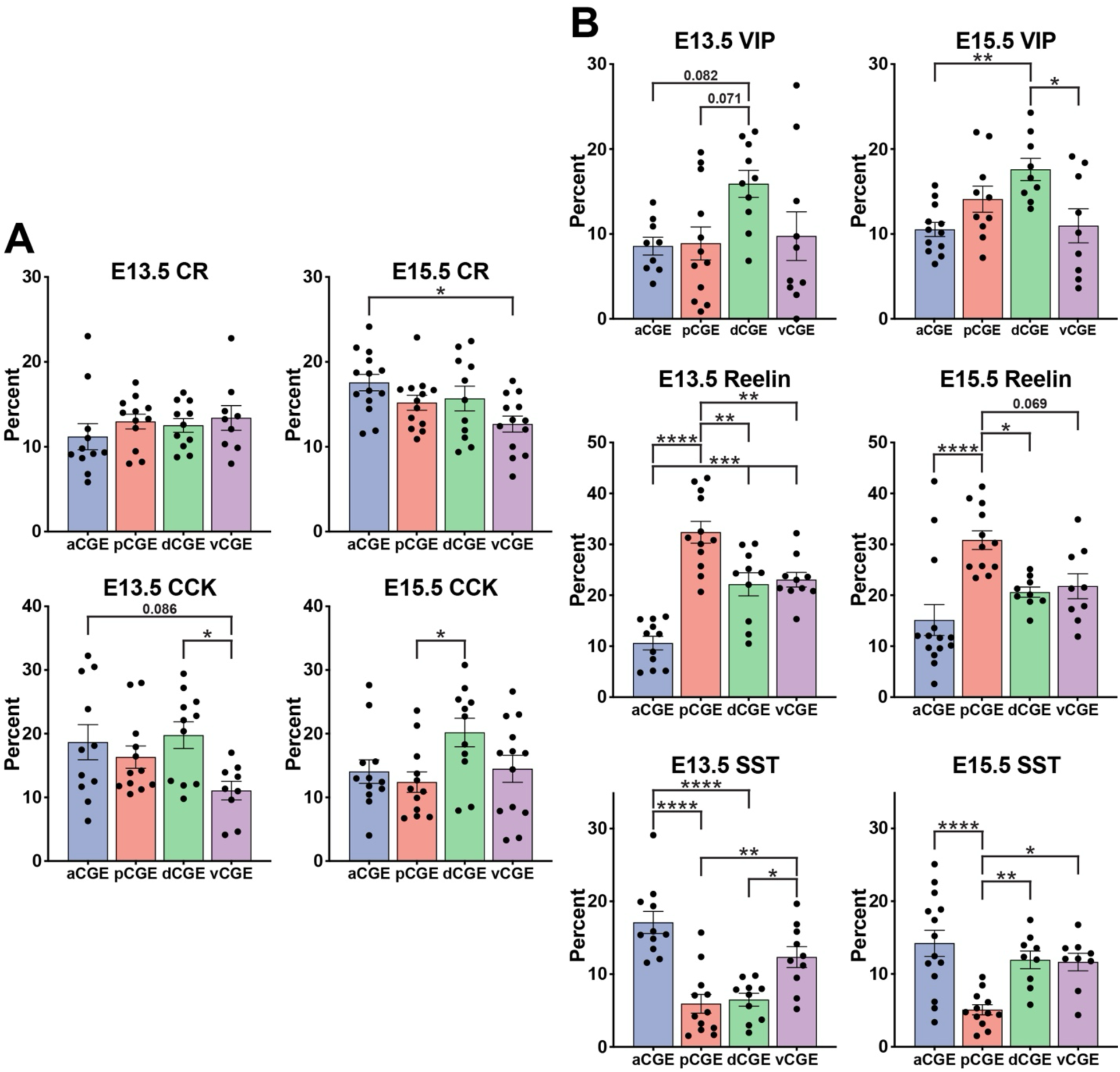
Significant biases for origins of interneuron subtypes from distinct CGE subdomains. Bar graphs displaying proportion of grafted Tom+ cells that express CCK, CR, VIP, Reln (Reln+/SST-) and SST (Reln+/SST+) from the 4 different CGE subdomains (aCGE, pCGE, dCGE, vCGE) at both E13.5 and E15.5 timepoints. * p < 0.05, ** p < 0.01, *** p < 0.0005, **** p < 0.0001.

VIP+ cells were enriched in the dCGE compared to other subdomains at both E13.5 and E15.5 (Figure 3B). At E13.5, ∼16% of Tom+ dCGE cells expressed VIP while it ranged from 8.6%-9.7% for the other 3 regions. Although this did not reach significance between any two CGE subdomains, there was a strong trend for increased VIP+ cells from dCGE compared to aCGE and pCGE (Figure 3B & Supplementary Table 2). At E15.5, there is a significant increase in VIP+ cells arising from dCGE compared to both aCGE and vCGE (Figure 3B & Supplementary Table 3).

Reln+ cells showed the strongest spatial bias of interneuron subtypes. The highest percentage of any subtype arising from any CGE subdomain was Reln+ cells from the pCGE; over 30% of pCGE Tom+ cells were Reln+ at both ages (Figures 3B & S4). Only 10.6% of E13.5 aCGE and 15.1% E15.5 aCGE cells were Reln+, while the dCGE and vCGE were between 20.6-23.0% at both ages; differences between these subdomains were significant for most of these comparisons (Figure 3B & Supplementary Tables 2-3). Thus, there is a clear spatial gradient of high Reln+ cells arising from the pCGE to fewer Reln+ cells arising from the aCGE at both embryonic ages.

Lastly, we observed spatial bias for SST+ cells that corresponds to the anatomical proximity of the MGE and CGE. At E13.5, there was a significant increase of SST+ cells in the aCGE and vCGE, with the pCGE having the fewest SST+ cells at both ages (Figure 3B & Supplementary Tables 2-3). The MGE merges into the CGE at the antero-ventral portion of the CGE, and expression of *Nkx2.1* extends towards the anatomical CGE in this region. Based on this relationship, we would expect the aCGE and vCGE to contain the greatest percentage of MGE-derived SST+ cells migrating caudally to the cortex.

### EdU labeling confirms biased origin of subtypes in CGE subdomains

To confirm that the relationship between spatial origin and mature CGE interneuron subtype arises from local neurogenesis in CGE subdomains, rather than migration of postmitotic cells through the CGE, we injected pregnant dams with EdU two hours prior to harvesting CGE tissue. This time frame should label ∼40-50% of actively dividing cells GE progenitors (52, 53) while minimizing the possibility of EdU-labeled cells migrating through the CGE and ‘contaminating’ the dissections.

We compared the percent of EdU+ cells in E15.5 transplants where we observed the largest spatial differences in subtypes: VIP+ cells between dCGE and aCGE, and Reln+ cells between pCGE and aCGE (Figures 3B & 4A). We also examined Reln+/SST+ cells in pCGE and aCGE to confirm that this population was EdU-negative and thus MGE-derived. As expected, almost no SST+ cells in the pCGE transplants were EdU+, while only ∼4.3% of SST+ cells in the aCGE were EdU+ (Figure 4B), confirming that these SST+ cells are indeed MGE-derived cells migrating through the CGE.

**Figure 4.**
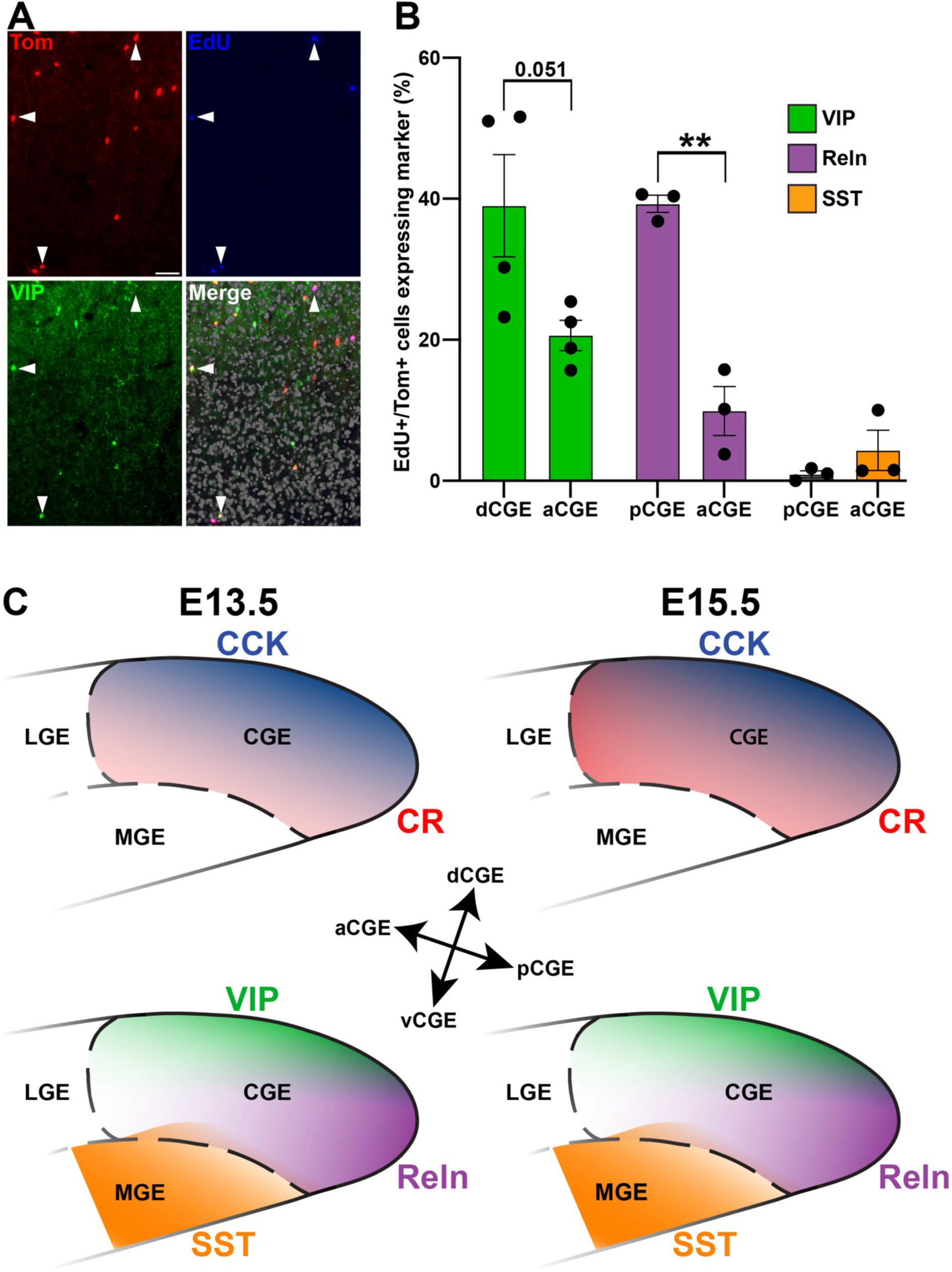
Significant biases for origins of interneuron subtypes from distinct CGE subdomains. **A.** Representative dCGE section showing several Tom+/EdU+ that co-express VIP (arrowheads). Scale bar = 50 μm. **B.** Graph depicting the percentage of Tom+/EdU+ cells that express VIP (dCGE & aCGE), Reln (pCGE & aCGE), and SST (pCGE & aCGE). ** p < 0.01. **C.** Schematic depicting enrichment of VIP+ and CCK+ cells in the dCGE, Reln+ cells in the pCGE, SST+ cells in the aCGE and vCGE, and an overall increase in CR+ cells at E15.5 compared to E13.5, with a significant increase in the aCGE.

Nearly 40% of all EdU+ cells in pCGE transplants were Reln+ while only ∼10% of EdU+ cells in aCGE transplants were Reln+ (Figure 4B). Although it did not reach statistical significance, a similar relationship exists for VIP+ cells, with 39.0% of EdU+ cells from dCGE transplants express VIP whereas only 20.6% of EdU+ cells from aCGE are VIP+ (p = 0.051) (Figure 4B). These findings confirm that there is a strong bias for Reln+ cells to originate from the pCGE and VIP+ cells to originate from the dCGE (Figure 4C).

## DISCUSSION

This study has defined a spatial and temporal map within the CGE relating mature interneuron subtypes with their spatial origin within the CGE (Figure 4C). We observed striking temporal consistency between E13.5 and E15.5 transplants, with the only significant difference being an overall increase in production of CR+ positive cells, and specifically from aCGE, at E15.5 (Figure 2C). This increased production of CR+ interneurons at E15.5 is consistent with a previous report (54), but that study compared E15.5 to E10.5-E12.5, timepoints when very few CGE cells are born (41). This finding is in sharp contrast to the MGE, where there is a clear relationship between temporal birthdate and the percentage of SST+ vs. PV+ cells that are generated (24). In fact, MGE-derived Chandelier cells are born from a small population of Nkx2.1+ progenitors during late embryogenesis (E15.5-E17.5), when the anatomical MGE no longer exists (19). It would be interesting to examine if this temporal consistency of CGE-derived interneurons is maintained at later embryonic timepoints, or if distinct subsets of these larger VIP+, CCK+, Reln+ and CR+ subtypes display biases in temporal birthdates.

While CR+ cells displayed almost no spatial bias, the other 3 CGE-derived interneuron subtypes displayed strong biases within the CGE. Reln+ cells were significantly enriched in the pCGE and displayed a low anterior-to-high posterior gradient. While VIP+ and CCK+ did not display an obvious gradient along an axis, both VIP+ and CCK+ cells were significantly enriched in dCGE transplants compared to other subdomains (Figures 3 & 4C). Several studies have argued that genetic signatures of mature interneuron subtypes only arise in postmitotic cells (55–57). However, the fact that EdU+ cells arising from distinct CGE subdomains show strong biases towards generating specific interneuron subtypes, as is the case in the MGE as well (17, 18), argues that critical fate-determining genetic programs exist in these cycling progenitors.

While transplant experiments have revealed critical insights into many aspects of neurodevelopment, they do circumvent some important developmental processes. For example, there is a period of programmed cell death from ∼P5-P12 in which ∼20-40% of cortical interneurons undergo apoptosis, which can be activity dependent or independent for different subtypes (58–62). Thus, in addition to intrinsic genetic programs, the final proportion of interneurons subtypes in brain regions can be sculpted by subtypes being preferentially susceptibility or resistant to programmed cell death. Whether programmed apoptosis occurs normally in our CGE transplants, or has differential effects on certain subtypes, is unclear, although there is evidence that apoptosis is intrinsically determined independent of various manipulations (58).

The E13.5 and E15.5 grafted CGE cells bypass their migration patterns to their proper cortical regions and instead are transplanted heterochronically into P1-P5 WT cortices. Despite these challenges, transplanted CGE cells still show a strong bias towards superficial cortical layers compared to MGE cells (Figure S1), mimicking their endogenous cortical patterning. While there is clear evidence that the age of host brains can strongly influence interneuron maturation, grafting cells into early postnatal brains yields superior survival and maturation (63). Additionally, the host brain environment (e.g., cortex vs. hippocampus) has a strong influence on the survival, maturation and relative proportion of interneuron subtypes in transplantation experiments (43, 63). Since CGE cells were grafted into the same WT brain regions for all transplants in this study, the local brain environment should not be a significant factor regulating differential interneuron survival from specific CGE subdomains. Last, there is evidence that distinct subtypes of MGE- and CGE-derived interneurons migrate through different streams to the cortex (64, 65). While different migratory streams can certainly provide critical information for targeting interneuron subtypes to specific brain regions, it is clear from many transplant studies that grafted interneurons mature and integrate into cortical circuitry while bypassing normal migratory routes.

The ability for grafted interneuron progenitors to integrate into local brain circuitry and inhibit hyperactive regions has shown great promise to treat a number of disorders such as epilepsy, schizophrenia, ASD and Alzheimer’s Disease (66, 67). While significant advances have been made to generate MGE-derived interneurons from human pluripotent cells for treating focal epilepsy (68, 69), differentiation protocols to obtain CGE-derived interneurons have not been established. Since dysfunction of CGE-derived interneurons has been associated with numerous neurodevelopmental disorders (36–38), linking this spatiotemporal map with differential gene expression within the CGE subdomains (40) will be critical for both (1) understanding genetic programs that drive CGE-derived interneurons, and (2) generating CGE-derived interneurons for therapeutic applications in the future.

## METHODS

### Animals

All experimental procedures were conducted in accordance with the National Institutes of Health guidelines and were approved by the NICHD Animal Care and Use Committee (protocol #23-047). *Dlx5/6-Cre* (Jax# 008199) (70), *Nkx2.1-Cre* (Jax# 008661) (71), and *Ai9* (Jax# 007909) (72) transgenic mouse lines were used in this study. For timed matings, noon on the day a vaginal plug was observed was denoted E0.5. Male and female embryonic donor mice and postnatal host mice were used without bias for all experiments.

### Harvesting embryonic CGE and MGE tissue

*Dlx5/6-Cre^C/+^*;*Ai9^F/+^* mice were used for all CGE dissections. E13.5- and E15.5-stage pregnant dams were injected with 5’-Ethunul-2’deoxyuridine (EdU, 100mg/kg) two hours prior to euthanasia. E13.5 and E15.5 embryonic brains were harvested as described previously (42). Briefly, pregnant dams were anesthetized via 5% isoflurane and euthanized via cervical dislocation. Embryos brains were removed and placed in sterile-filtered ice cold carbogenated artificial cerebral spinal fluid (ACSF, in mM: 87 NaCl, 26 NaHCO_3_, 2.5 KCl, 1.25 NaH_2_PO_4_, 0.5 CaCl_2_, 7 MgCl_2_, 10 glucose, 75 sucrose, saturated with 95% O_2_, 5% CO_2_, pH 7.4), and checked under a fluorescent scope to confirm tdTomato expression. For each embryo, the MGE was removed and then the LGE-CGE continuum was hemisected to separate the LGE and CGE. To collect aCGE and pCGE, the CGE was divided into three equivalent pieces, and the anterior and posterior regions were collected (middle third discarded). To collect dCGE and vCGE, the CGE was hemisected longitudinally and the dorsal and ventral halves were collected. Process repeated for all embryos, and each CGE region from 1-2 litters (4-10 embryos depending on experiment) were pooled together in ice cold ACSF in 5-mL round bottom tubes prior to preparation of single cell suspensions.

*Nkx2.1-Cre^C/+^*;*Ai9^F/+^* mice were used for MGE dissections. Whole MGE from E13.5 embryos were dissected and pooled together in ice cold ACSF in 5-mL round bottom tubes prior to preparation of single cell suspensions.

### Preparing single cell suspensions

Single cell suspensions were generated as previously described (42–44). ACSF was removed from each tube, replaced with 1 mL of freshly prepared Pronase solution (1 mg/mL Pronase in ACSF) and incubated at RT for 15-20 minutes. Pronase solution was removed and replaced with 1 mL of reconstitution solution (1% Fetal Bovine Serum + DNase I (2U/μL) in ACSF). Tissue was mechanically dissociated via trituration with fire-polished glass Pasteur pipettes, first with large bore opening (∼500 μm) followed by smaller bore (∼100-200 μm). Single cell suspensions were then passed through 35 μm filter on a 5 mL round bottom tube (Corning# 352235) pre-wetted with ACSF. Solution was transferred to 1.5 mL Protein LoBind tubes (Eppendorf# 022431081) and spun down in a swinging bucket centrifuge at 500 g for 5 minutes. Supernatant was removed, leaving ∼20-40 μL of cell suspension. Cells were carefully resuspended and counted on a Countess 3 Cell counter. Cell suspensions were diluted in reconstitution solution as needed to generate single cell suspension of ∼10,000-25,000 cells/μL, depending on total cells recovered and number of pups to inject.

### Cell transplantations

Cell transplantations were performed as previously described (43, 44). P2-P5 postnatal WT pups were anesthetized on ice for 4-5 minutes, then secured on a petri dish to stabilize the head and visualize lambda. Angled pulled glass micropipettes filled with mineral oil were loaded into a Nanoject III Programmable Nanoliter Injector. Single cell suspensions were front loaded into the micropipette and the Nanoject head secured onto a micromanipulator. Micropipette tip was centered perpendicular over lambda. Each pup received 4 cortical injections, 2 in each hemisphere with the following coordinates from lambda: 1.0 mm anterior, 1.0 mm lateral, 0.75-1.0 mm depth; and 2.0 mm anterior, 1.0 mm lateral, 0.75-1.0 mm depth. Nanoject injection conditions at each site: 10 pulses of 60 nL each, injection rate of 30, 1 second delay between each pulse. After injection, pups are placed on heating pad until fully recovered from anesthesia, then returned to their nest.

### Brain harvesting and immunohistochemistry

30-35 days post transplantation, mice were anesthetized with Euthasol (270 mg/kg) and transcardially perfused with 4% paraformaldehyde (PFA) in PBS. Brains were removed and post-fixed in 4% PFA for 0-2 hours, then transferred to PBS. Brains were sectioned at 50 μm on a vibratome and stored as floating sections in antifreeze solution (30% ethylene glycol, 30% glycerol, 40% PBS) at −20°C in 96-well plates. Free floating sections from were washed in PBS to remove antifreeze solution and incubated in blocking solution (10% Normal Donkey Serum in PBS + 0.3% Triton X-100) for 1-2 hours. Sections were incubated with primary antibodies in blocking solution for 48-72 hours at 4°C, then washed in PBS for 2-4 hours at RT. Sections were incubated with secondary antibodies with DAPI in blocking solution O/N at 4°C, washed in PBS, mounted and imaged at 20X on a Zeiss Axioimager.M2 with ApoTom.2 (with Zen Blue software) or an Olympus VS200 Slide Scanner (VS200 ASW).

The following antibodies were used in this study: rat-anti SST (1:300, Millipore MAB354), rabbit anti-VIP (1:500, ImmunoStar 20077), rabbit anti-proCCK (1:1000, Frontier Institute MSFR105030), mouse anti-Reelin (1:1000, Sigma-Aldrich MAB5364), mouse anti-Calretinin (1:1000, Millipore MAB1568), goat-anti PV (1:1000, Swant PVG-213). CCK & CR were co-stained in one set of sections while VIP, Reln and SST were co-stained in another set of sections. Species-specific fluorescent secondary antibodies used were conjugated to AlexaFluor^®^ 488, 647 and 790, all used at 1:500. For some experiments, brain sections were stained for EdU expression using the Click-IT EdU Cell Proliferation Kit (Invitrogen# C10337) after immunostaining. Images were processed with Adobe Photoshop and/or Fiji.

### Cell Counting

All CGE cell counts were performed by hand and blind to genotypes. Total brains counted for each condition are as follows: E13.5, aCGE = 9-11 brains from 7 experiments, pCGE = 12 brains from 5 experiments, dCGE = 10-11 brains from 4 experiments, vCGE = 9 brains from 4 experiments; E15.5, aCGE = 12-14 brains from 5 experiments, pCGE = 10-13 brains from 5 experiments, dCGE = 9-11 brains from 4 experiments, vCGE = 9-13 brains from 4 experiments. A total of 99 mice were used for all conditions. Range of brains is due to different brain numbers used for distinct immunostaining conditions. Only Tom+ cells that had migrated away from the injection site were counted, and any Tom+ cells located outside of cortical layers I-VI were excluded. Range of Tom+ cells counted per brain per immunostaining condition was 97-546, with an average of 410 Tom+ cells counted per brain for all experimental and immunostaining conditions combined.

For EdU cell counts, a minimum of 50 EdU+/Tom+ cells were counted per brain, with n = 3-4 brains per CGE transplant condition for VIP, Reln and SST immunostaining. Range of EdU+/Tom+ cells counted per brain per immunostaining condition was 56-109, with an average of 77 EdU+ cells counted per brain. We estimate that ∼10-20% of all Tom+ cells co-expressed EdU in each brain.

Imaris (Oxford Instruments, v10.2) was used to determine the relative distance of grafted Tom+ MGE and pCGE cells between the pial surface and white matter (WM) border below layer VI. We chose pCGE since it had the lowest percentage of migrating MGE cells in the CGE. The ‘Spots’ tool enabled automated cell detection, with manually defined thresholds refined using Tom+/DAPI+ cells as a reference, and non-cells were manually removed. The ‘Surface’ tool was used to generate borders along the pial surface and the WM. ‘Object-Object Statistics’ yielded the shortest distance of each cell to both borders. Distance was normalized between images by using image voxel size from VS200 ASW (x=0.325μm/pixel; y=0.325μm/pixel). This data set was exported to Excel and each cell’s cortical depth was determined with the following calculation:

(Distance to WM / (Distance to Pia + Distance to WM)) x 100 In this equation, a cell on the pia surface would have a score of 100 and a cell at the deep WM border would have a score of 0. Cells were also grouped into 10 equidistant bins spanning the cortical thickness, with scores 100-90 in bin 1, 90-80 in bin 2, etc., allowing for calculation of the percent of cells in each bin (i.e., cortical depth) for each brain. n = 5 MGE brains, range of 134-556 Tom+ cells/brain; n = 4 pCGE brains, range of 115-251 Tom+ cells/brain.

### Statistics

All cell counts were collected in Excel and analyzed with Prism (v10.4.2). Ordinary one-way ANOVA was performed for all analysis of > 2 datapoints, followed by Tukey’s multiple comparison t-test to determine significance between all datapoints. Unpaired t-tests were used to determine significance between E13.5 and E15.5 for each cell type and brain region, and for differences in cortical depth between CGE and MGE transplants. All statistics (F and P values for ANOVAs, P values for t-tests) are presented in Supplementary Tables 1-3.

## ACKNOWLEDGEMENTS

We thank members of the Petros lab for comments on this project and manuscript; W. Emrick for some cell counts; M. Anath & M. Gastinger for help with the Imaris analysis.

## FUNDING & CONFLICTS OF INTEREST

This work was supported by *Eunice Kennedy Shriver* NICHD Intramural award to T.P. The authors declare that the research was conducted in the absence of any conflict of interest.

---

**Figure S1.**
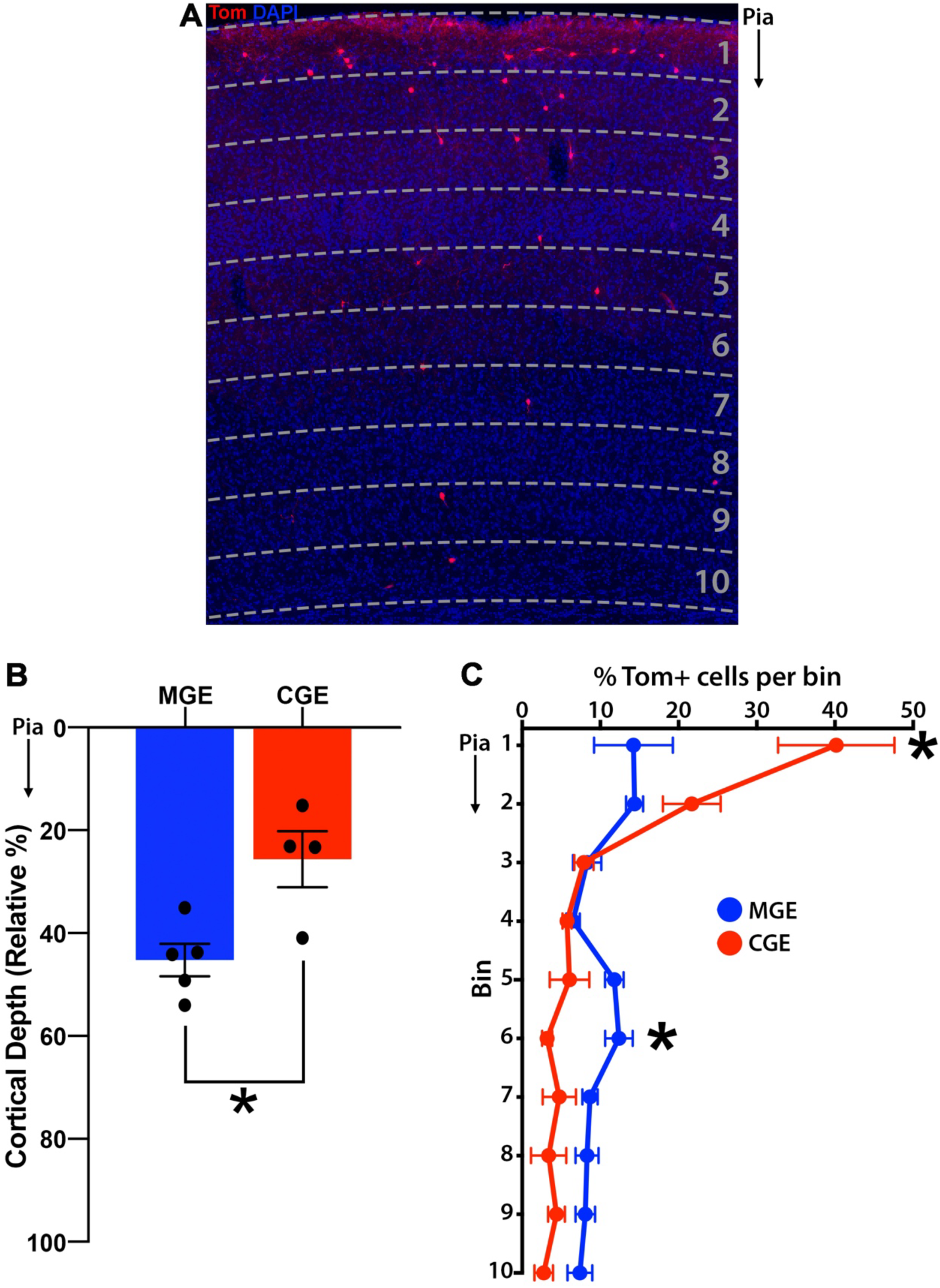
Transplanted CGE-derived cells are biased towards superficial layers. **A.** Example of grafted Tom+ cells with equidistant bins spanning from the pial surface to the layer VI boundary. **B.** Bar graph depicting average depth of grafted Tom+ MGE and CGE cells, ranging from 0 at the pia surface to 100 at the layer VI-subplate boundary. * p < 0.05. **C.** Line graph depicting the percent of grafted Tom+ MGE and CGE cells in 10 equivalent size bins spanning from the pia (Bin 1) to the white matter boundary below layer VI (Bin 10).

**Figure S2.**
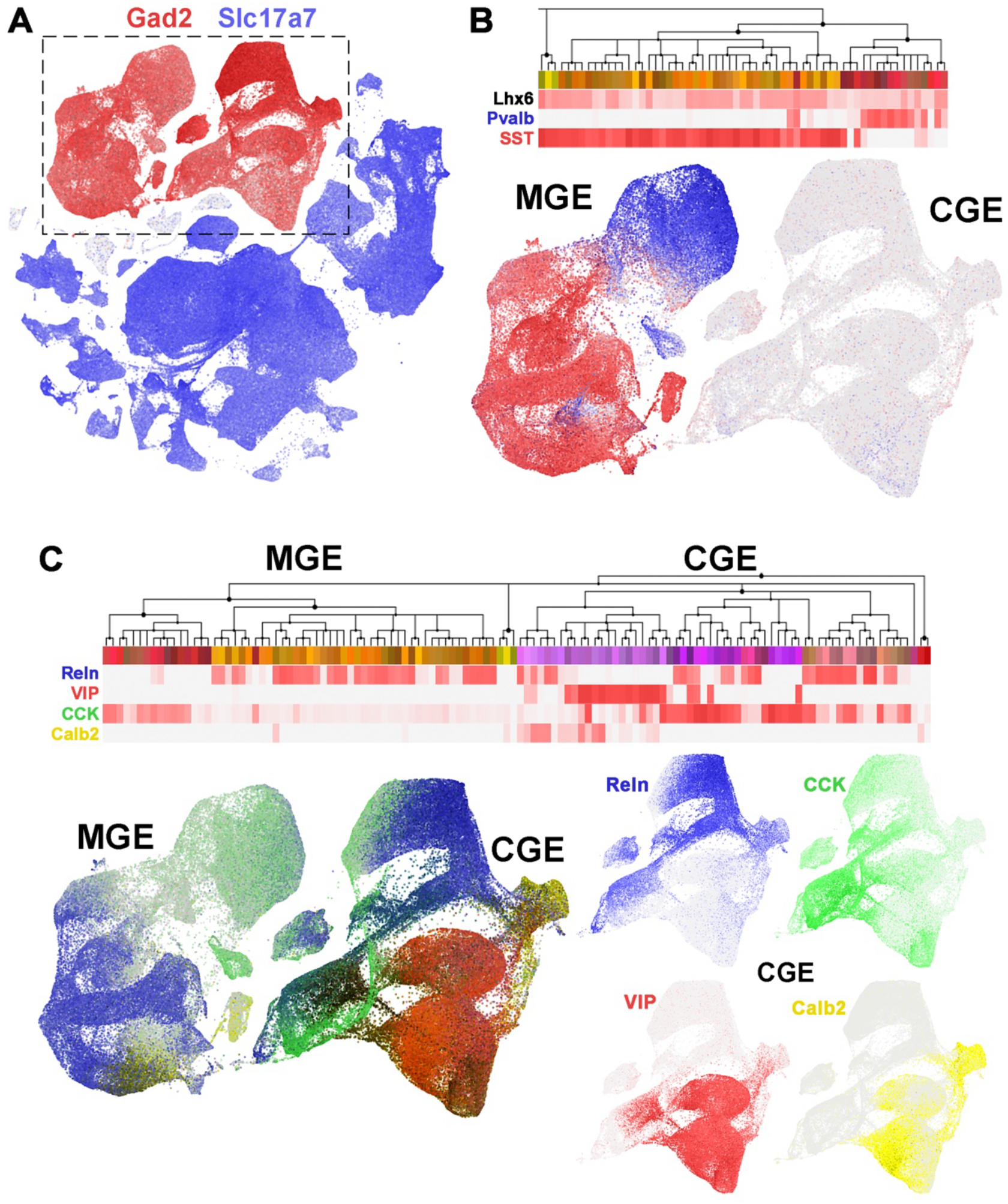
Markers of mature CGE-derived interneuron subtypes. All heatmaps and scatterplots are taken and adapted from the Allen Brain Map Cell Types Database: RNA-Seq Data (portal.brain-map.org/atlases-and-data/rnaseq), specifically the mouse whole cortex & hippocampus –10x Genomics with 10x smart-sequencing taxonomy (47). **A.** Scatterplot of all cells in the dataset; inhibitory and excitatory neurons can be cleanly segregated by expression of Gad2 and Slc17a7, respectively. **B.** Heatmap (top) and scatterplot (bottom) demonstrating MGE-derived interneurons can be cleanly segregated into 2 cardinal subclasses expressing either Pvalb (PV) or SST. **C.** Heatmap (top) and scatterplot (bottom) showing expression of 4 markers used in this study to define CGE-derived interneurons, highlighting the complimentary expression patterns of VIP (red) and Reln (blue), and CCK (green) and CR (yellow).

**Figure S3.**
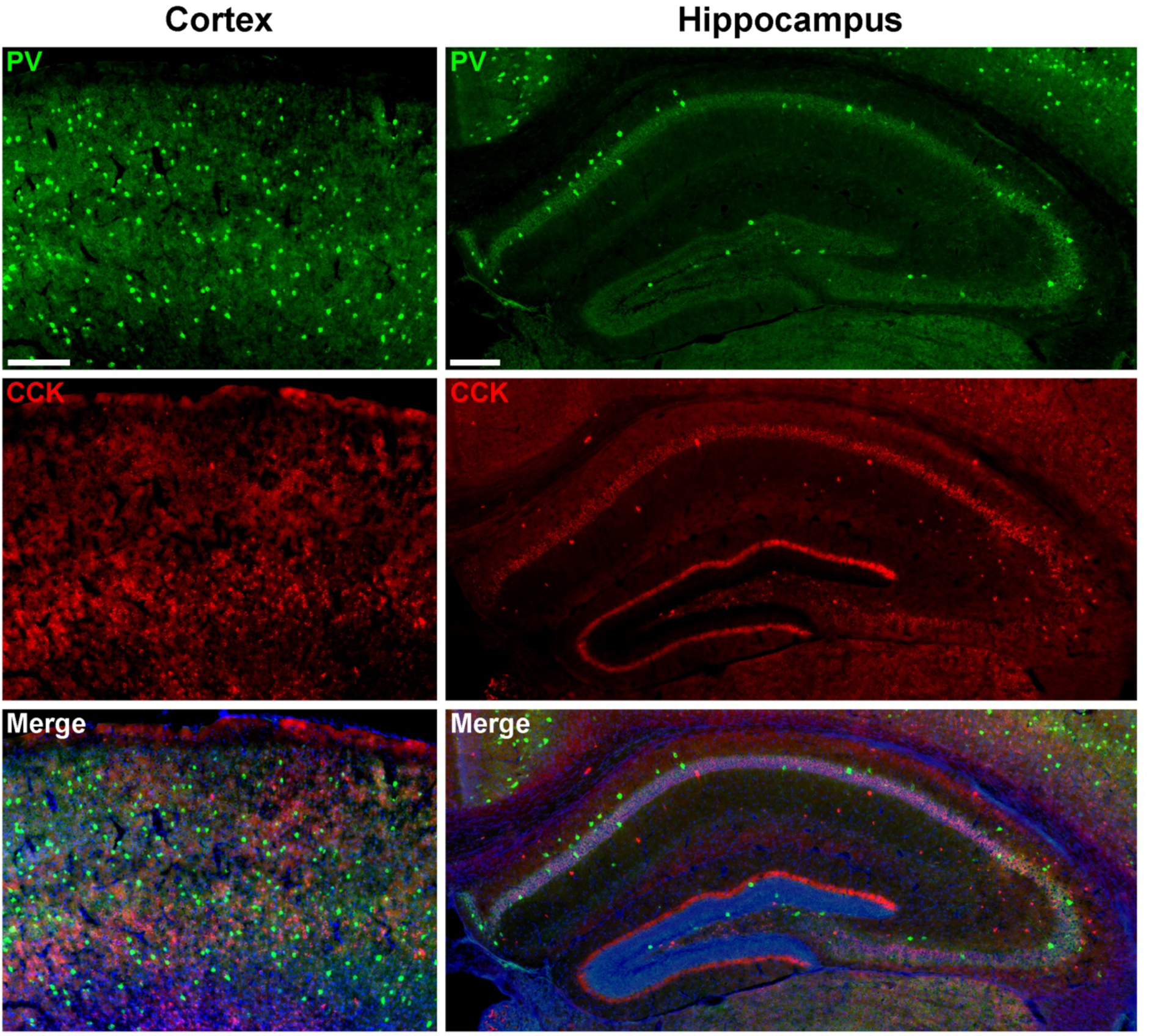
MGE-derived PV+ interneurons do not express CCK protein. Representative images through the cortex and hippocampus showing lack of colocalization between PV and CCK proteins. Scale bars = 100 μm.

**Figure S4.**
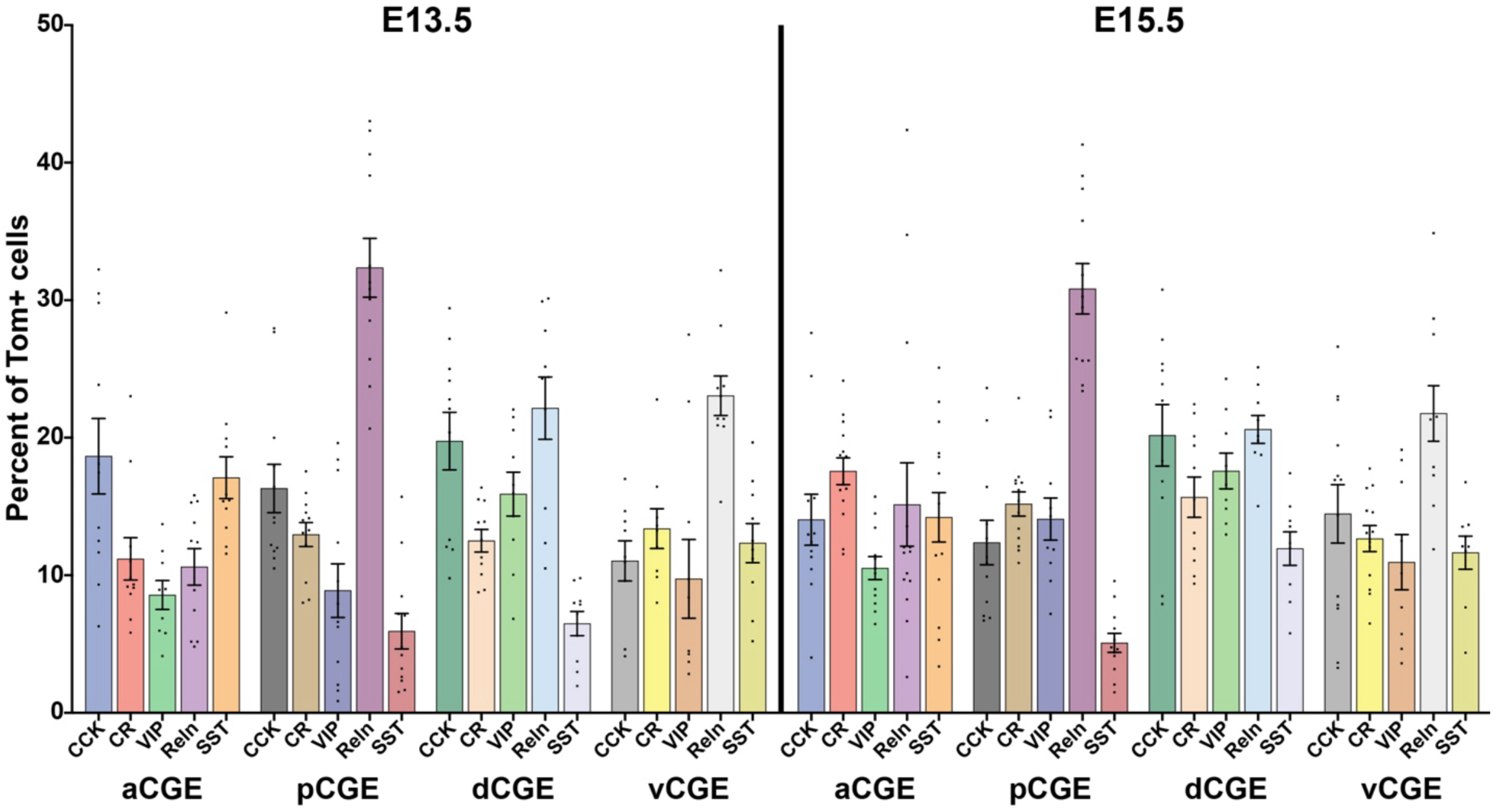
All datapoints used for analysis. Graph depicting all datapoints for each immunostained cell maker CCK, CR, VIP, Reln (Reln+/SST-) and SST (Reln+/SST+) for all 8 experimental conditions; 4 CGE subdomains at E13.5 (left) and E15.5 (right).

## SUPPLEMENTARY TABLE LEGENDS

**Supplementary Table 1.** Statistics for E13.5 vs. E15.5 timepoints. Unpaired t-tests were used to compare E13.5 vs. E15.5 for all CGE regions for all cell types, and for all regions combined for each cell type (last column). P-values presented for each t-test. Significant results are bold, results with a trend that did not reach significance (0.10 > p > 0.05) are highlighted in red. ** p < 0.01.

| **E13.5 vs.**<br>**E15.5** | **aCGE** | **pCGE** | **dCGE** | **vCGE** | **All regions**<br>**combined** |
| --- | --- | --- | --- | --- | --- |
| **CCK** | NS<br>0.172 | NS<br>0.113 | NS<br>0.892 | NS<br>0.243 | NS<br>0.317 |
| **CR** | **<br>**0.001** | NS<br>**0.087** | NS<br>**0.073** | NS<br>0.666 | **<br>**0.001** |
| **VIP** | NS<br>0.156 | NS<br>**0.055** | NS<br>0.430 | NS<br>0.739 | NS<br>0.123 |
| **ReIn+/SST-** | NS<br>0.224 | NS<br>0.593 | NS<br>0.556 | NS<br>0.650 | NS<br>0.866 |
| **ReIn+/SST+** | NS<br>0.249 | NS<br>0.570 | **<br>**0.002** | NS<br>0.716 | NS<br>0.801 |

**Supplementary Table 2.** Statistics for all E13.5 transplants. One-way ANOVA was used to compare all 4 brain regions, followed by Tukey’s multiple comparison t-test between 2 regions. F and P values given for each ANOVA (right column), and adjusted P-value presented for each t-test. Significant results are bold, results with a trend that did not reach significance (0.10 > p > 0.05) are highlighted in red. * p < 0.05, ** p < 0.01, *** p < 0.0005, **** p < 0.0001.

| **E13.5** | **aCGE**<br>**vs.**<br>**pCGE** | **aCGE**<br>**vs.**<br>**dCGE** | **aCGE**<br>**vs.**<br>**vCGE** | **pCGE**<br>**vs.**<br>**dCGE** | **pCGE**<br>**vs.**<br>**vCGE** | **dCGE**<br>**vs.**<br>**vCGE** | **ANOVA comparing**<br>**all 4 regions** |
| --- | --- | --- | --- | --- | --- | --- | --- |
| **CCK** | NS<br>0.850 | NS<br>0.982 | NS<br>**0.086** | NS<br>0.636 | NS<br>0.327 | **\***<br>**0.039** | **\***<br>**P value = 0.042**<br>F = 3.008 |
| **CR** | NS<br>0.691 | NS<br>0.855 | NS<br>0.590 | NS<br>0.992 | NS<br>0.995 | NS<br>0.957 | NS<br>P value = 0.600<br>F = 0.637 |
| **VIP** | NS<br>0.999 | NS<br>**0.082** | NS<br>0.979 | NS<br>**0.071** | NS<br>0.990 | NS<br>0.162 | **\***<br>**P value = 0.049**<br>F = 2.874 |
| **ReIn+/SST-** | **\*\*\*\***<br>**<0.0001** | **\*\*\***<br>**0.0005** | **\*\*\***<br>**0.0002** | **\*\***<br>**0.0019** | **\*\***<br>**0.0051** | NS<br>0.987 | **\*\*\*\***<br>**P value < 0.0001**<br>F = 24.52 |
| **ReIn+/SST+** | **\*\*\*\***<br>**<0.0001** | **\*\*\*\***<br>**<0.0001** | NS<br>**0.072** | NS<br>0.990 | **\*\***<br>**0.007** | **\***<br>**0.021** | **\*\*\*\***<br>**P value < 0.0001**<br>F = 16.57 |

**Supplementary Table 3.** Statistics for all E15.5 transplants. One-way ANOVA was used to compare all 4 brain regions, followed by Tukey’s multiple comparison t-test between 2 regions. F and P values given for each ANOVA (right column), and adjusted P-value presented for each t-test. Significant results are bold, results with a trend that did not reach significance (0.10 > p > 0.05) are highlighted in red. * p < 0.05, ** p < 0.01, *** p < 0.0005, **** p < 0.0001.

| **E15.5** | **aCGE**<br>**vs.**<br>**pCGE** | **aCGE**<br>**vs.**<br>**dCGE** | **aCGE**<br>**vs.**<br>**vCGE** | **pCGE**<br>**vs.**<br>**dCGE** | **pCGE**<br>**vs.**<br>**vCGE** | **dCGE**<br>**vs.**<br>**vCGE** | **ANOVA comparing**<br>**all 4 regions** |
| --- | --- | --- | --- | --- | --- | --- | --- |
| **CCK** | NS<br>0.933 | NS<br>0.152 | NS<br>0.999 | *<br>**0.043** | NS<br>0.870 | NS<br>0.189 | NS<br>**P value = 0.051**<br>F = 2.797 |
| **CR** | NS<br>0.367 | NS<br>0.604 | **<br>**0.008** | NS<br>0.989 | NS<br>0.333 | NS<br>0.221 | *<br>**P value = 0.015**<br>F = 3.842 |
| **VIP** | NS<br>0.260 | **<br>**0.005** | NS<br>0.996 | NS<br>0.336 | NS<br>0.431 | *<br>**0.016** | **<br>**P value = 0.004**<br>F = 5.226 |
| **ReIn+/SST-** | ****<br>**<0.0001** | NS<br>0.401 | NS<br>0.237 | *<br>**0.032** | NS<br>**0.069** | NS<br>0.990 | ***<br>**P value = 0.0002**<br>F = 8.203 |
| **ReIn+/SST+** | ****<br>**<0.0001** | NS<br>0.663 | NS<br>0.570 | **<br>**0.009** | *<br>**0.014** | NS<br>0.999 | ***<br>**P value = 0.0001**<br>F = 8.852 |

## Notes

### Competing Interest Statement

The authors have declared no competing interest.

